# HIV-1 requires capsid remodelling at the nuclear pore for nuclear entry and integration

**DOI:** 10.1101/2021.03.18.436028

**Authors:** Anabel Guedán, Callum D Donaldson, Ophélie Cosnefroy, Ian A Taylor, Kate N Bishop

**Affiliations:** Retroviral Replication Laboratory, The Francis Crick Institute, London, United Kingdom; Macromolecular Structure Laboratory, The Francis Crick Institute, London, United Kingdom

**Author notes:** These authors contributed equally (Anabel Guedán, Callum D Donaldson).

## Abstract

The capsid (CA) lattice of the HIV-1 core plays a key role during infection. From the moment the core is released into the cytoplasm, it interacts with a range of cellular factors that, ultimately, direct the pre-integration complex to the integration site. For integration to occur, the CA lattice must disassemble. Early uncoating or a failure to do so has detrimental effects on virus infectivity, indicating that an optimal stability of the viral core is crucial for infection. Here, we introduced cysteine residues into HIV-1 CA in order to induce disulphide bond formation and engineer hyper-stable mutants that are slower or unable to uncoat, and then followed their replication. From a panel of mutants, we identified three with increased capsid stability in cells and found that, whilst the M68C/E212C mutant had a 5-fold reduction in reverse transcription, two mutants, A14C/E45C and E180C, were able to reverse transcribe to approximately WT levels. Moreover, these mutants only had a 5-fold reduction in 2-LTR circle production, suggesting that not only could reverse transcription complete in hyper-stable cores, but that the nascent viral cDNA could enter the nuclear compartment. Furthermore, we observed significant levels of A14C/E45C mutant capsid in nuclear and chromatin-associated fractions implying that the hyper-stable cores themselves entered the nucleus. Immunofluorescence studies revealed that although the A14C/E45C mutant capsid reached the nuclear pore with the same kinetics as wild type capsid, it was then retained at the pore in association with Nup153. Crucially, infection with the hyper-stable mutants did not promote CPSF6 re-localisation to nuclear speckles, despite the mutant capsids being competent for CPSF6 binding. These observations suggest that hyper-stable cores are not able to uncoat, or remodel, enough to pass through or dissociate from the nuclear pore and integrate successfully. This, is turn, highlights the importance of capsid lattice flexibility for nuclear entry. In conclusion, we hypothesise that during a productive infection, a capsid remodelling step takes place at the nuclear pore that releases the core complex from Nup153, and relays it to CPSF6, which then localises it to chromatin ready for integration.

**AUTHOR SUMMARY:** The mature viral core of human immunodeficiency virus (HIV) consists of a highly organised lattice formed by capsid molecules that encloses the viral RNA and viral enzymes. This lattice is crucial during the early stages of viral replication, as it has to break down – uncoat – at the right time and place in order for the viral DNA to integrate successfully. Lentiviruses, like HIV, can infect non-dividing cells and are able to access the host cell DNA by entering the nucleus through nuclear pores. Until recently, uncoating was thought to occur in the cytoplasm as the whole core was thought too large to pass through the nuclear pore. However, lately it has been suggested that uncoating might occur at the nuclear pore or even inside the nucleus and the site of uncoating is currently hotly debated. By investigating HIV mutants with an increased lattice stability, we have shown that lattice flexibility is crucial for nuclear entry. Provocatively, we observed hyper-stable mutant capsid in nuclear and chromatin-associated fractions suggesting that uncoating is not required for nuclear entry. Nonetheless, microscopy experiments suggested that these hyper-stable mutants were retained on the inner side of the nuclear pore, and were impaired for downstream events in the nucleus, leading to a severe infectivity defect. Therefore, we believe that an essential uncoating, or capsid lattice remodelling event normally takes place at the nuclear pore.

## INTRODUCTION

Upon infection of the target cell, the human immunodeficiency virus 1 (HIV-1) core is released into the cytoplasm before trafficking to the nucleus. During this process, the viral RNA genome is reverse transcribed by the viral reverse transcriptase (RT) into double-stranded DNA forming what is known as the reverse transcription complex (RTC). Inside the nucleus, the viral DNA is the main component of the pre-integration complex (PIC) and is integrated by the viral integrase (IN) into the host cell genome to form a provirus [1].

The mature core of HIV-1 consists of a highly organised lattice of capsid (CA) molecules, encasing the viral RNA genome and associated viral proteins. The lattice is composed of approximately 1500 CA monomers assembled into about 250 hexamers and exactly 12 pentamers, forming a fullerene cone shape [2–5]. The CA monomer is a largely helical protein with two independent folded domains separated by a flexible linker: the N-terminal domain (NTD) and the C-terminal domain (CTD) [2, 6–8]. Structural studies have provided key information on how the CA lattice is organised [8–14]. Intra-hexamer and inter-hexamer interactions at different lattice interfaces contribute to an optimal CA lattice stability. The intra-hexamer interactions include NTD-NTD and NTD-CTD interactions between adjacent CA monomers within a hexamer, whereas the inter-hexamer interactions include dimeric and trimeric CTD-CTD interactions between neighbouring hexamers [11, 14].

As the outer surface of the viral core, the CA lattice protects the viral DNA from cytosolic sensors [15, 16] but it can be recognised by cellular restriction factors that inhibit viral replication [17, 18]. Moreover, from cell entry to integration, the CA lattice interacts with many cellular factors. Some of these are cytoplasmic, including Cyclophilin A (CypA) [19–21], the cellular motors dynein and kinesin-1 [1, 22, 23] and FEZ1 [24]. Others shuttle between the cytoplasm and the nucleus like Transportin-3 (TNPO3) [25–27] and Transportin-1 [28]. There are further interactions at the nuclear envelope with nuclear pore complex (NPC) proteins, particularly nucleoporins 358 (Nup358) and 153 (Nup153) [25, 26, 29–31] and with nuclear proteins such as cleavage and polyadenylation factor 6 (CPSF6); [31–35]. The roles of most of these cellular factors in viral replication are not yet fully understood.

Prior to integration, the CA lattice is assumed to disassemble in a process termed uncoating; however, the timing, location and even the exact definition of this process remains unclear. For many years, uncoating was believed to occur immediately after viral entry [36, 37] but, more recently, data has suggested that it might occur after a short time in the cytoplasm, at the nuclear pore or even, most recently, inside the nucleus [38–44]. Indeed, there is growing evidence to suggest that at least some CA is present in the nucleus, although its oligomeric state and function have yet to be fully characterised [34, 40–43, 45–48]. For this reason, using the term “remodelling” may be more appropriate than uncoating to suggest changes in the CA lattice. We, and others, have also reported a link between CA loss and reverse transcription [38, 49–51]. Uncoating is inhibited if reverse transcription is stalled. Initially, this was taken as evidence for cytoplasmic uncoating [49]. However, when combined with emerging data suggesting CA can reach the nucleus more rapidly than previously thought [43, 44, 48], it has reopened the debate about where reverse transcription is actually completed during infection, and so does not currently preclude any specific uncoating model.

Importantly, the location and timing of uncoating together with optimal stability of the CA lattice seem to be key to successful infection. Early uncoating, or a failure to uncoat, both have detrimental effects on viral infectivity [15, 52, 53], making the CA lattice an attractive target for drug development [54, 55]. Here we set out to investigate the effects of increased core stability on the early stages of HIV-1 infection. To this end, we introduced mutations into CA predicted to stabilise the mature CA lattice at different interfaces. We show that mutations that create a hyper-stable lattice reduce virus infectivity by inhibiting integration, but only slightly impede reverse transcription. Analysis of CA protein levels within different subcellular fractions during infection revealed higher levels of hyper-stable mutant CA in all fractions over time, including in the nucleus. Finally, immunofluorescence data suggest that the hyper-stable mutant CA lattice is retained around the nuclear pore and that it is unable to promote CPSF6 re-localisation to nuclear speckles. Based upon these observations, we propose a model where hyper-stable mutants are unable to uncoat or “remodel” their capsid lattice to the required extent to successfully deliver the viral DNA for integration into the host cell genome.

## RESULTS

### Cysteine mutations at different CA lattice interfaces are able to stabilise the viral core in cells

Previous *in vitro* studies introduced cysteine mutations at the CA NTD-NTD interface to create disulphide bridges in order to stabilise CA hexamers for crystallisation [11, 12]. We decided to use the same approach to examine if similar mutations would increase the stability of the CA lattice during infection of cells, to investigate the effect of core stability on the early stages of HIV-1 infection. We selected CA mutants from the literature and identified additional CA residues to substitute with cysteine residues in order to stabilise all of the different inter-hexamer interfaces with disulphide bridges (listed in Fig 1A and residues highlighted in Fig 1B). The new mutants E180C, L151C/L189C and K203C/A217C were designed based on a previous report demonstrating that disulphide bonds can have a variable Cβ-Cβ inter-residue spacing of between 3.5-4.5Å [56]. Thus, employing a cryo-EM MDFF atomic model of an *in vitro* CA tubular assembly [PDB ID: 3J34; [8]] and crystal structures [PDB ID: 3H4E and 3H47; [11]], we selected residue pairs with a Cβ-Cβ distance within this range at the various interfaces for site-directed mutagenesis. Consequently, we created a panel with at least two mutants at each CA lattice interface. In addition, the P38A mutant was included in the panel as a negative control, as it has shown to be less stable than WT [51, 52]. The original mutant used to determine the crystal structure of a hexameric CA assembly, A14C/E45C/W184A/M185A, [11, 12] was also included as a negative control.

**Figure 1.**
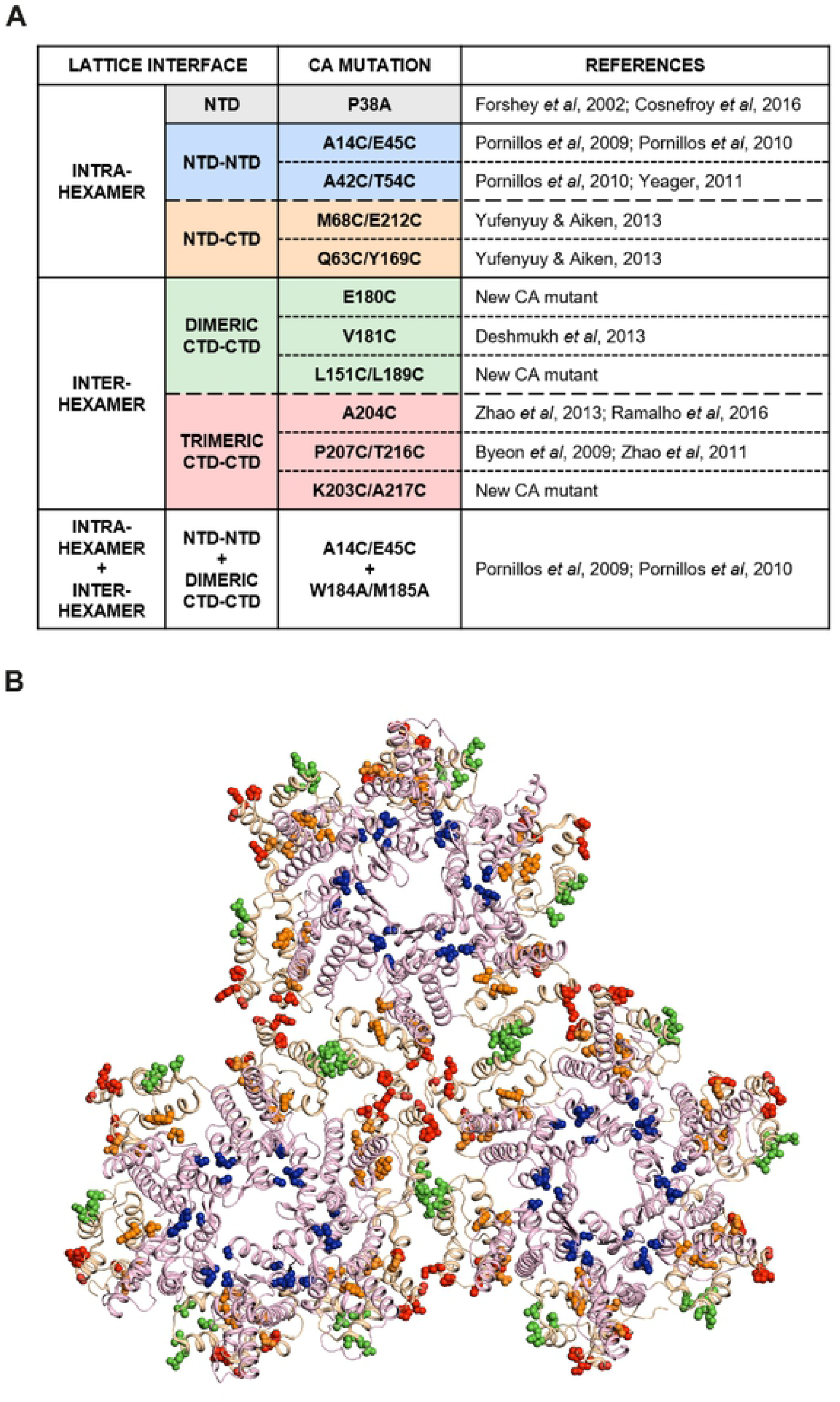
Panel of CA mutants. (A) The panel of CA mutants used in this study and the lattice interface at which the mutations reside. New CA mutants with cysteine mutations (E180C, L151C/L189C and K203C/A217C) were designed based on a previously published cryo-EM MDFF atomic model (PDB ID: 3J34; Zhao et al, 2013) and crystal structures (PDB ID: 3H4E and 3H47; Pornillos et al, 2009). The remaining CA mutants were selected from previous publications as indicated in the references column. (B) Structure of the CA lattice from PDB ID: 3J34, showing 18 CA monomers arranged into three hexameric rings. Residues where mutations were made are highlighted and colour-coded according to the lattice interface type, as in (A).

Firstly, we synthesised viral-like particles (VLP) expressing the different CA mutants and a GFP reporter gene. We assessed VLP production by measuring the RT activity in the supernatants of producer 293T cells using a modified RT ELISA. As expected, the A14C/E45C/W184A/M185A and W184A/M185A mutants that weaken CTD-CTD interactions were severely impaired for virus production (Supplementary Fig 1A). We confirmed that this was not due to a lack of Gag expression of these mutants, by immunoblotting 293T producer cell lysates with an anti-HIV-1 CA antibody and showing that the level of Gag expression in the mutants was similar to WT (Supplementary Fig 1B). Most of the mutants, however, showed similar titres to WT VLP, implying that these mutations did not affect Gag assembly. Of note, the inter-hexamer mutants V181C, L151C/L189C and K203C/A217C were partially impaired for VLP production (Supplementary Fig 1A), likely due to the importance of this interface on virus assembly [3, 53]. Whilst we confirmed that expression of these mutant capsid proteins in 293T producer cell lysates was similar to WT VLP (Supplementary Fig 1C), we detected an additional, smaller, CA band for both L151C/L169C and K203/A217C. This has been reported previously [12, 57] and probably represents a processing defect, such as inappropriate cleavage of CA by the viral protease due to the structural changes [57]. This may be the cause of the reduced viral titres. Thus, based on these results, the L151C/L189C and the K203C/A217C mutants as well as the A14C/E45C/W184A/M185A and W184A/M185A mutants were not included in further experiments.

Next, we performed fate-of-capsid assays, to study the effect of the cysteine mutations on viral core stability in cells (Fig 2). HeLa cells were infected with equal titres of WT and mutant VLP, and cell lysates were harvested at 2 and 20 hours post-infection (hpi). These time points were selected as they have previously been used to monitor the stability of the viral core at the early and late stages of the virus replication cycle [17]. The cell lysates were centrifuged through a sucrose cushion to be separated into two fractions: free or soluble CA (S) and pelleted CA (P), which contains assembled cores [58]. An input (I) sample was also harvested before centrifugation through the sucrose cushion. The fractions were analysed by immunoblotting with an anti-HIV-1 CA antibody and the ratio of soluble to pelleted CA was determined. Fig 2 shows representative blots of the fate-of-capsid assay for each mutant. A WT sample was included in every assay for comparison. Although there was some variation between assays, the WT samples showed similar amounts of CA in the pellet and soluble fractions (P=S) at 2 hpi and less CA in the pellet (P<S) at 20hpi (Fig 2A), presumably reflecting a reduction in the amount of assembled CA with time due to uncoating. In agreement with previous reports, we recovered much less CA in the pellet fraction than the supernatant fraction from cells infected with the negative hypo-stable control, P38A, [51, 52]. For this mutant, P<S at both 2 and 20 hpi, indicating the small amount of assembled CA at both time points. Therefore, we set a CA ratiometric profile criteria for mutants to be considered hyper-stable as the following: Mutants should consistently show greater amounts of CA in the pellet than in the supernatant (P>S) at 2 hpi and at least equal amounts in both fractions (P=S) at 20 hpi. Three mutants showed this phenotype: M68C/E212C, A14C/E45C and E180C (highlighted with dashed boxes in Fig 2A). The profile of P207C/T216C and the V181C mutants was similar to WT (Fig 2A). At 2 hpi, A42C/T54C showed a CA ratio P=S and, Q63C/Y169C and A204C showed P<S, suggesting that none of these mutants had increased viral core stability compared to WT. Thus, taken together despite the cytoplasm being a reducing environment [59, 60], these data showed that three of the panel of cysteine mutants had increased core stability in cells and could be used to study the effects on replication of stabilizing the viral core.

**Figure 2.**
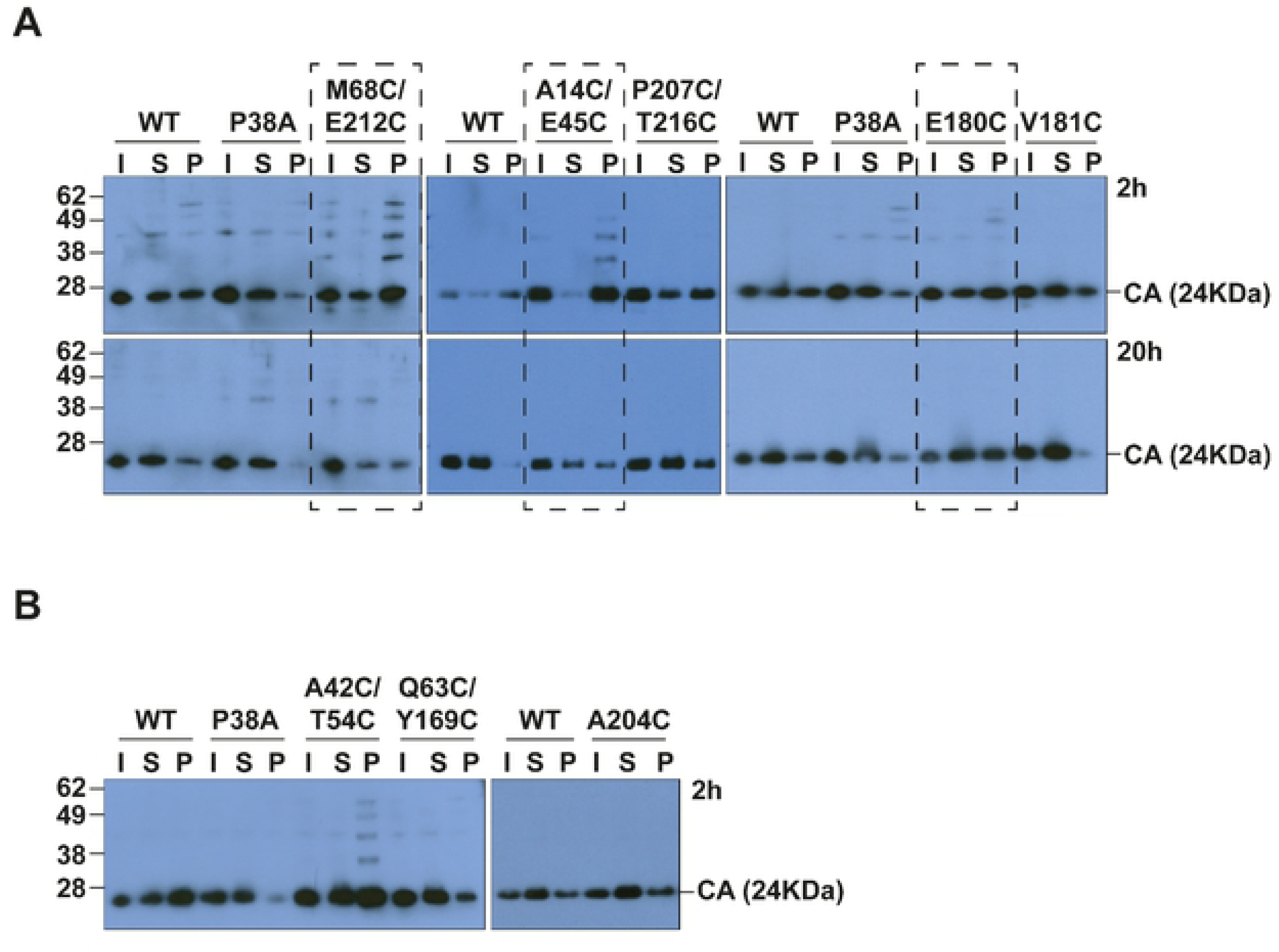
Effect of the CA mutations on viral core stability in cells. (A, B) Representative immunoblots of Fate-of-capsid assays comparing WT VLP to mutant VLP. HeLa cells were infected with equal RT units of WT or mutant VLP and cell lysates were harvested at 2 hpi (A, B) and 20 hpi (A). Cell lysates were centrifuged through a sucrose cushion to separate viral CA into free (soluble, S) and assembled (pellet, P) fractions. An input (I) sample was also harvested before the centrifugation through the sucrose cushion. CA was detected by immunoblotting using an anti-HIV-1 CA antibody. Mutant VLP with a CA ratio of P>S at 2h and P=S at 20h were considered “hyper-stable” (surrounded with a dashed-line box). Each assay was performed at least three times independently.

### Most CA mutants are less infectious, regardless of CA lattice stability

To investigate the effect of the CA mutations on virus infectivity, we infected four cell lines (293T, HeLa, SupT1 and U937 cells) with equal RT units of VSVg-pseudo-typed GFP-reporter WT or mutant VLP and analysed the percentage of GFP+ cells at 72 hpi by flow cytometry (Fig 3). As observed previously, the P38A mutant that had reduced CA lattice stability had a marked reduction in GFP expression in all the cell lines (Fig 3, grey bars). Apart from the inter-hexamer P207C/T216C mutant that showed similar infectivity to WT, the remainder of the mutants also showed decreased infectivity, ranging from ∼0.05-10% of WT infectivity. A similar pattern of infectivity was seen across all the cells lines indicating that there were no cell-type specific effects, with generally lower overall infectivity in U937 cells. Interestingly, the level of infectivity did not correlate with the CA lattice stability determined by the fate-of-CA assay (Fig 2) but there was some correlation with the CA interface modified (see colour coding Fig 1A and Fig 3), with the intra-hexamer mutants (blue and orange bars) being more defective than the inter-hexamer mutants (green and red bars). Of the three hyper-stable mutants, A14C/E45C, M68C/E212C and E180C (Fig 3, black arrowheads), the infectivity ranged from 0.7-4%, 0.07-0.4% and 2-10% respectively between the different cell lines.

**Figure 3.**
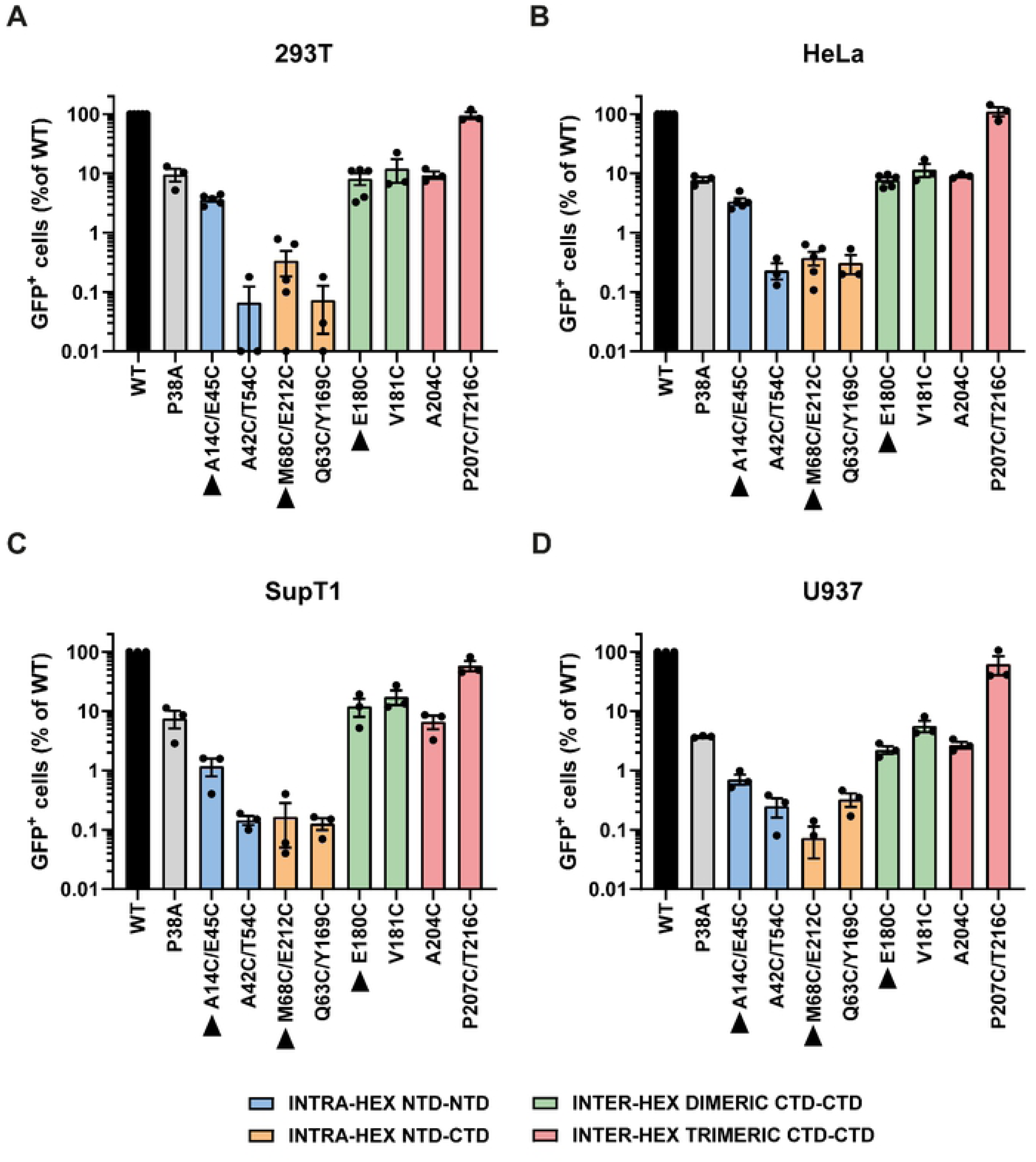
Effect of CA mutations on VLP infectivity. 293T (A), HeLa (B), SupT1 (C) and U937 (D) cells were infected with equal RT units of GFP-reporter WT or mutant VLP. The percentage of GFP+ cells was measured by flow cytometry at 72hpi and plotted relative to WT VLP. Points indicate individual biological repeats and lines show the mean ± SEM. Bars are colour coded according to the lattice interface at which the cysteines have been introduced, as in Fig 1. Hyper-stable mutants based on the fate of capsid assay are indicated with black arrow heads.

### Reverse transcription can complete in hyper-stable cores but infectivity is severely inhibited

Next, to determine which step in replication was affected by the CA mutations, we examined the ability of each mutant to reverse transcribe (Fig 4). 293T cells were synchronously infected with equal RT units of WT or mutant VLP, and at 0, 1, 2, 4, 6 and 24 hpi, cells were harvested and the DNA extracted and analysed for early (strong stop) and late (second strand) viral cDNA products by qPCR (Fig 4). Following infection with WT VLP, the amount of viral cDNA products increased with time, peaking at 6 hpi (Fig 4A-D, black line). As seen before, infection with mutant P38A resulted in less accumulation of viral cDNA (Fig 4A, grey lines). This ∼90% decrease in cDNA accumulation mirrored the 90% decrease in infectivity (Fig 3). Likewise, the P207C/T216C mutant had similar infectivity to WT VLP and had similar levels of reverse transcription (Fig 4B, red lines). However, the hyper-stable mutants A14C/E45C and M68C/E212C had a different phenotype. Surprisingly, despite a ∼95% decrease in infectivity, A14C/E45C reverse transcribed to WT levels (Fig 4C, blue lines) and mutant M68C/E212C still produced about 10% the cDNA of WT (Fig 4D, orange lines), even though its infectivity was reduced to less than 1% (Fig 3). Figures 4E and 4F show a summary of the amount of strong stop cDNA accumulated at 6 and 24 hpi, respectively, for all the mutants. The E180C and V181C mutants showed WT levels of strong stop cDNA, A204C accumulated about 10% the cDNA of WT and A42C/T54C and Q63C/Y169C only accumulated ∼1% the cDNA of WT. Similar results were obtained when looking at the levels of second strand cDNA (Fig S2A). In general, the level of cDNA accumulation at 24 hours compared to WT correlated with relative level of infectivity compared to WT (Fig 4G and Fig S2B). Plotting these data as a ratio of relative cDNA levels to relative infectivity shows that most mutants have a ratio of less than 3 (Fig 4H and Fig S2C). Therefore, it can be assumed that, in general, the reduction in infectivity results from the effect on reverse transcription. The only mutants where the level of reverse transcription products did not correlate with the level of infectivity was for the three hyper-stable mutants (Fig 4H and Fig S2C). These data suggest that even though increasing core stability has detrimental effects on infectivity, the block is after reverse transcription. This implies that the core does not need to uncoat in order to reverse transcribe. Therefore, although reverse transcription promotes uncoating of WT virus [38, 49–51, 61], reverse transcription is not dependant on uncoating.

**Figure 4.**
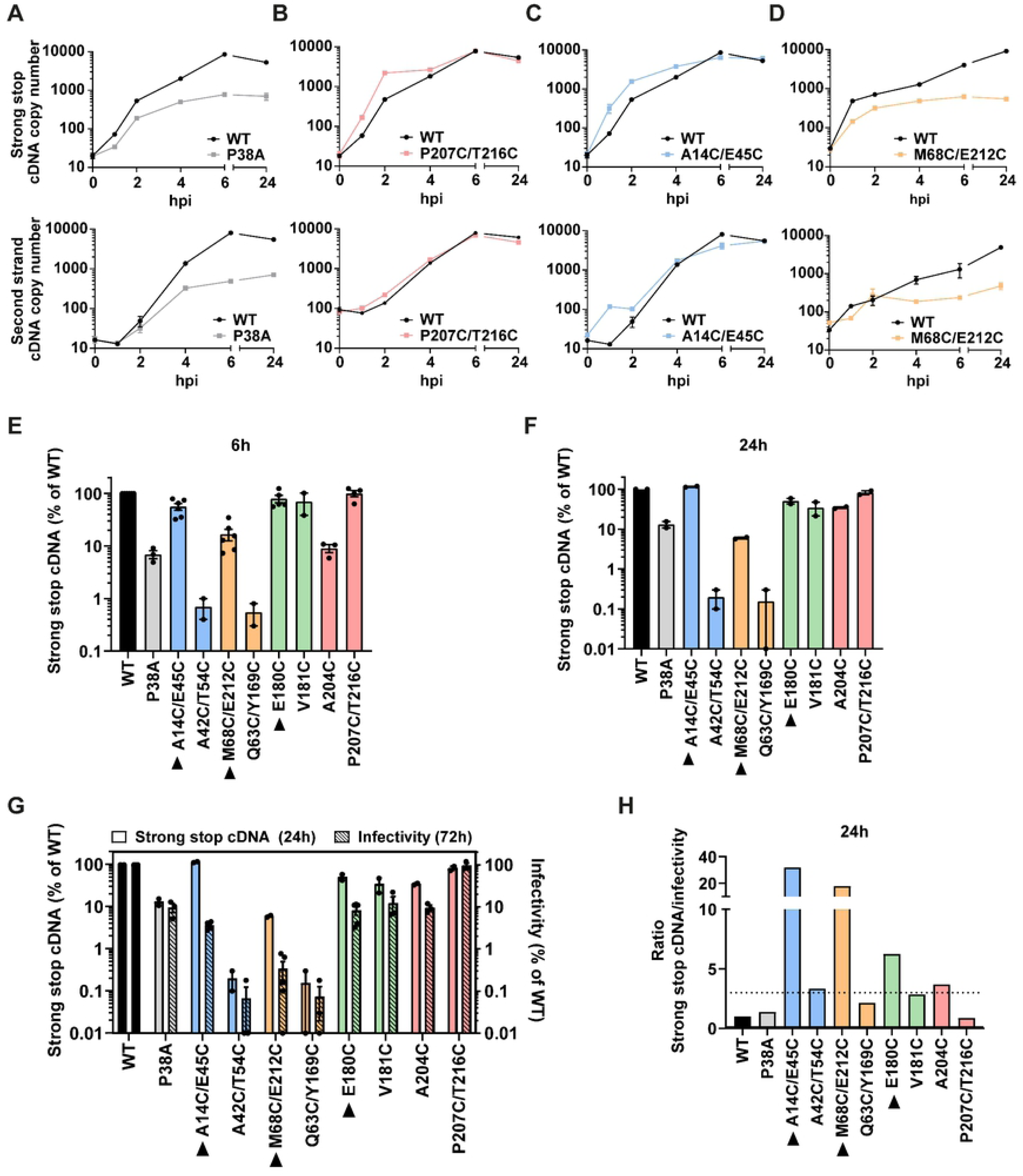
Effect of CA mutations on reverse transcription. 293T cells were synchronously infected with equivalent RT units of WT or mutant VLP. At the indicated times post-infection, cells were harvested for DNA extraction, and viral cDNA products were measured by qPCR. (A-D) Graphs show the levels of early (strong stop) cDNA (upper panels) and late (second strand) cDNA (lower panels) in 293T cells following infection with WT VLP (Black line) and mutants P38A (A), P207C/T216C (B), A14C/E45C (C) or M68C/E212C (D). The data is shown as mean ± SEM from at least two independent experiments. (E, F) Bar charts show the levels of strong stop cDNA at 6h (E) and 24h (F) post infection for the panel of mutants relative to WT infections. (G) Bar chart showing the levels of strong stop cDNA at 24 h (left y-axis) and infectivity (from Fig 3) at 72 h (right y-axis) compared to WT VLP for each mutant. Individual points represent biological repeats and lines indicate the mean +/-SEM. (H) Bar chart showing the ratio of relative levels of strong stop cDNA to infectivity, from (G). Dashed line indicates a ratio of 3. Bars are colour coded according to the lattice interface at which the cysteines have been introduced, as in Fig 1. Hyper-stable mutants are indicated with a black arrow heads.

We therefore decided to focus on the A14C/E45C, E180C and M68C/E212C mutants to investigate the effect of core stability on the other early replication steps. First, we examined whether the observed increase in core stability of these mutants was indeed due to disulphide bridge formation as designed, and not due to other stabilising effects of the individual mutations. HeLa cells were infected with equal RT units of WT and mutant VLP and cell lysates were analysed by SDS-PAGE in reducing or non-reducing conditions followed by immunoblotting with an anti-HIV-1 CA antibody (Fig 5A). Samples were either treated with iodoacetamide to prevent artefactual cysteine formation (non-reducing conditions), or dithiothreitol (DTT) to reduce disulphide bonds (reducing conditions) prior to SDS-PAGE. Only monomeric CA of 24KDa was detected for WT VLP, in both reducing and non-reducing conditions. By contrast, disulphide cross-linking of CA monomers was detected in the non-reducing conditions for all three hyper-stable mutants. A strong band was evident at 48 kDa, which corresponds to a CA dimer, and slower migrating species were also present, suggesting higher order oligomers existed. Upon addition of DTT, the higher molecular weight bands disappeared, suggesting that intra- or inter-hexamer disulphide bonds contribute to the increased core stability of these mutants.

**Figure 5.**
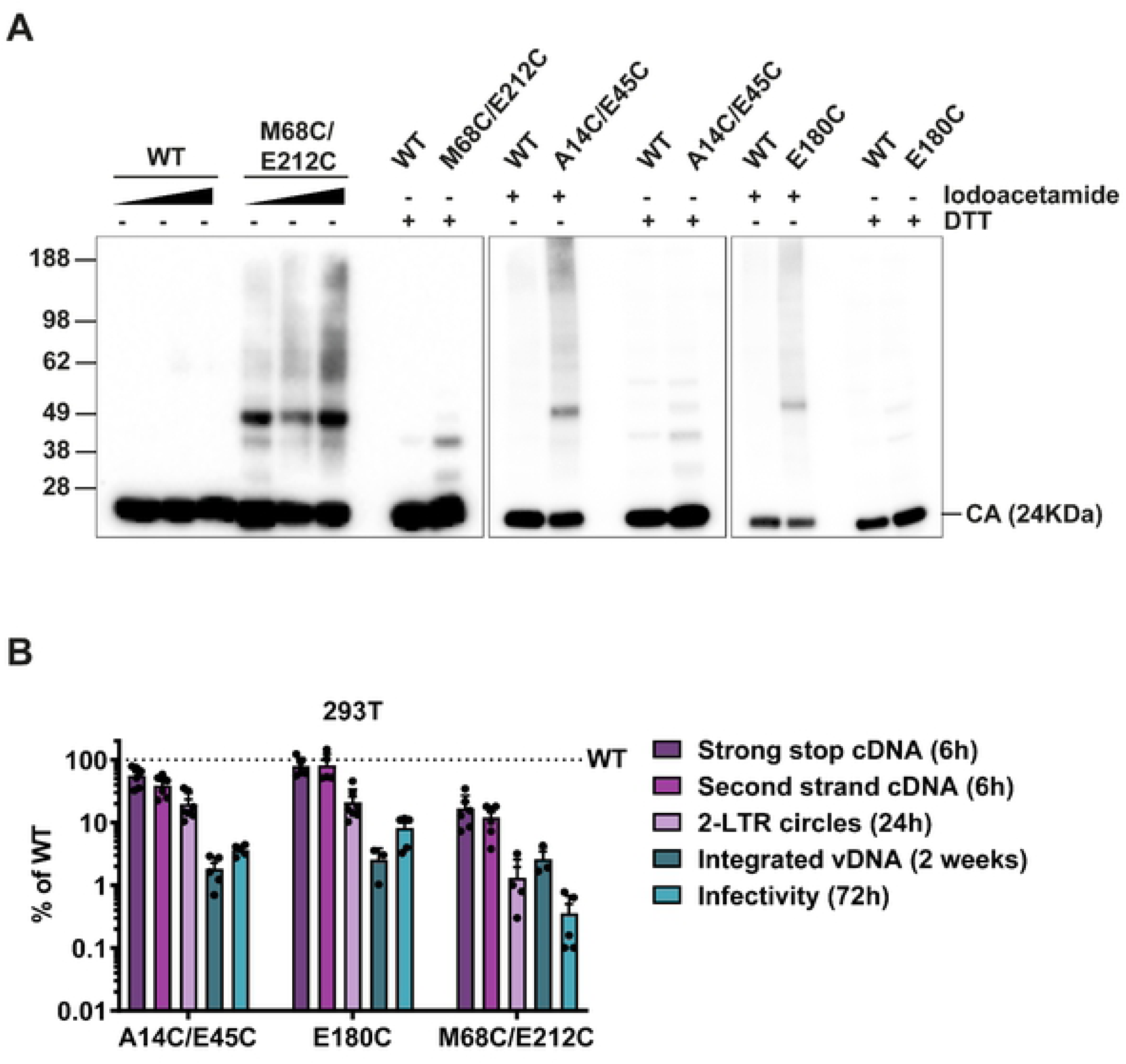
Effect of hyper-stable CA mutations on the early stages of infection. (A) Disulphide cross-linking of CA monomers in cells. HeLa cells were infected with WT VLPs or the hyper-stable mutants M68C/E212C, A14C/E45C or E180C, and cell lysates were analysed by non-reducing SDS-PAGE and immunoblotting with an HIV-1 CA antibody. Samples were either treated with 10µM-50µM (indicated by gradient arrows in first panel) or 50µM (indicated by “+”) Iodoacetamide, to prevent further disulphide bond formation, or with 0.1M DTT (indicated by “+”), to reduce existing disulphide bonds, prior to SDS-PAGE. The prominent bands at 24 and 48 kDa correspond to monomeric and dimeric CA, respectively. (B) 293T cells were synchronously infected with equivalent RT units of WT or mutant VLP. At the indicated times post-infection, cells were harvested for DNA extraction, and early and late cDNA (6hpi), 2-LTR circles (24hpi) and integrated proviral DNA (2 weeks post infection) were measured by qPCR. Data were plotted relative to WT infections (shown as a dashed line at 100%). Points indicate individual biological repeats and lines show the mean ± SEM. Viral infectivity from parallel infections (see Fig 3) was plotted for comparison.

To further investigate the infectivity block to the hyper-stable mutants, we measured the ability of the mutants to complete different steps of replication between reverse transcription and integration. Following synchronous infection of 293T cells with equal RT units of WT or mutant VLP, samples were taken at 6 hpi, 24 hpi and two weeks post infection, and levels of early and late cDNA, 2-LTR circles and integrated viral DNA were quantified by qPCR. It is widely accepted that 2-LTR circles are an indicator of completion of reverse transcription and nuclear entry. Fig 5B shows the relative amounts of the different viral DNA products compared to WT VLP. Interestingly, as well as being only slightly defective for reverse transcription, both A14C/E45C and E180C mutant VLP were able to produce 2-LTR circles at ∼20% the amount of 2-LTR circles produced by WT VLP. This suggests that at least some of these mutant particles can enter the nucleus. Importantly, these mutants integrated <2% of the amount of cDNA integrated in WT infections, which correlated well with the decrease in infectivity measured. Therefore, there appears to be a minor block to nuclear entry for these two mutants, but, surprisingly, the major defect is after nuclear entry and before integration. In contrast, mutant M68C/E212C VLP only produced 10% of the cDNA of WT, and there was a further reduction in the amount of 2-LTR circles, to <1% of that produced by WT VLP. There was no further decrease in integration compared to the levels of 2-LTR circles, suggesting that the M68C/E212C mutant is more severely impaired earlier in replication than the other two hyper-stable mutants and is blocked for nuclear entry. Altogether, these data suggests that reverse transcription can finish in a hyper-stable core and that this cDNA is able to enter the nuclear compartment, but that there is a block to integration that prevents these hyper-stable mutants successfully forming proviruses.

### CA protein from the A14C/E45C mutant is detected in nuclear and chromatin fractions

The nuclear pore has generally been considered too small to allow passage of a complete HIV core, and yet we have shown here that hyper-stable CA mutants can synthesise cDNA and that at least some of this cDNA can reach the nucleus to form 2-LTR circles. This suggests that either the cDNA can escape the core in order to enter the nucleus, despite the lattice being more resistant to disassembly, or that the whole stable lattice can enter the nucleus. Therefore, in order to determine the cellular localisation of components of a hyper-stable core, we measured CA and IN protein levels in different cellular compartments during infection (Fig 6). HeLa cells were synchronously infected with equal RT units of WT or A14C/E45C VLP and cells were harvested at 4, 8, 24 and 30 hpi. The total levels of both CA and IN in whole cell lysates were similar for WT and A14C/E45C (CC) infection at all time points, both decreasing between 8 and 24 hpi (Fig 6A). The infected cell lysates were next processed into the following subcellular fractions: cytoplasm, membranes, nucleus and chromatin-bound. The separation resolution of the subcellular fractionation was determined by looking for distinct markers in the different fractions: HSP90 for the cytoplasm, Calnexin for the membranes, HDAC2 for the nucleus and Histone H3 for chromatin-bound. The representative immunoblot in Fig 6B shows that the fractionation procedure was highly effective, and the markers were predominantly found in the expected fractions. We also probed for Tubulin-α and Lamin B1 to monitor the distribution of the cytoskeleton and the nuclear envelope, respectively. Tubulin-α was highly enriched in the cytoplasmic fraction while Lamin B1 was mainly localised in the nuclear fraction and a small portion seemed to be chromatin-bound. Furthermore, we confirmed that CPSF6 is mainly found in the soluble nuclear fraction. Having determined that the cellular compartments had been separated successfully, we analysed CA and IN protein levels in the different subcellular fractions by immunoblotting with anti-HIV-1 CA and anti-HIV-1 IN antibodies. The amount of CA in the cytoplasm remained relatively constant throughout the course of the infection (Fig 6C), with noticeably more CA detected following infection with A14C/E45C VLP, especially at 8 and 24 hpi. Conversely, IN levels decreased markedly between 8 and 24 hpi, but there was little difference between WT and A14C/E45C infections (Fig 6C). A similar pattern was observed in the membrane fraction (Fig 6D). This confirms the increased stability of the A14C/E45C CA lattice compared to WT CA, and suggests that IN protein has a shorter half-life than CA. Surprisingly, we detected both CA and IN in nuclear fractions as early as 4 hpi (Fig 6E). Particularly surprising was the amount of A14C/E45C CA detected. Also unexpected was that more CA and IN were detected in the nucleus at the earlier time points than at 24 hours. This suggested that viral cores travel to the nucleus faster than previously thought, and before the peak of reverse transcription at 6 hpi (see Fig 4). Other groups have also recently reported this phenomenon [39, 40, 43, 44, 48]. Finally, we observed a similar, but more extreme, pattern of detection of CA in the chromatin fractions (Fig 6F). Here we observed significantly more A14C/E45C CA than WT CA present at all time points, and, again, A14C/E45C CA was present event at 4 hpi. Interestingly, there was more IN present in this fraction from the WT infections than the A14C/E45C infections at all time points. The association of A14C/E45C CA with nuclear fractions, particularly with the chromatin-bound fraction, suggests that, despite being hyper-stable, this CA protein could enter the nucleus. However, as the nuclear envelope marker, Lamin B1, was also distributed in these two fractions, it could also be hypothesised that A14C/E45C was accumulating at the nuclear envelope rather than inside the nucleus. To help resolve these options, we performed cellular localisation experiments.

**Figure 6.**
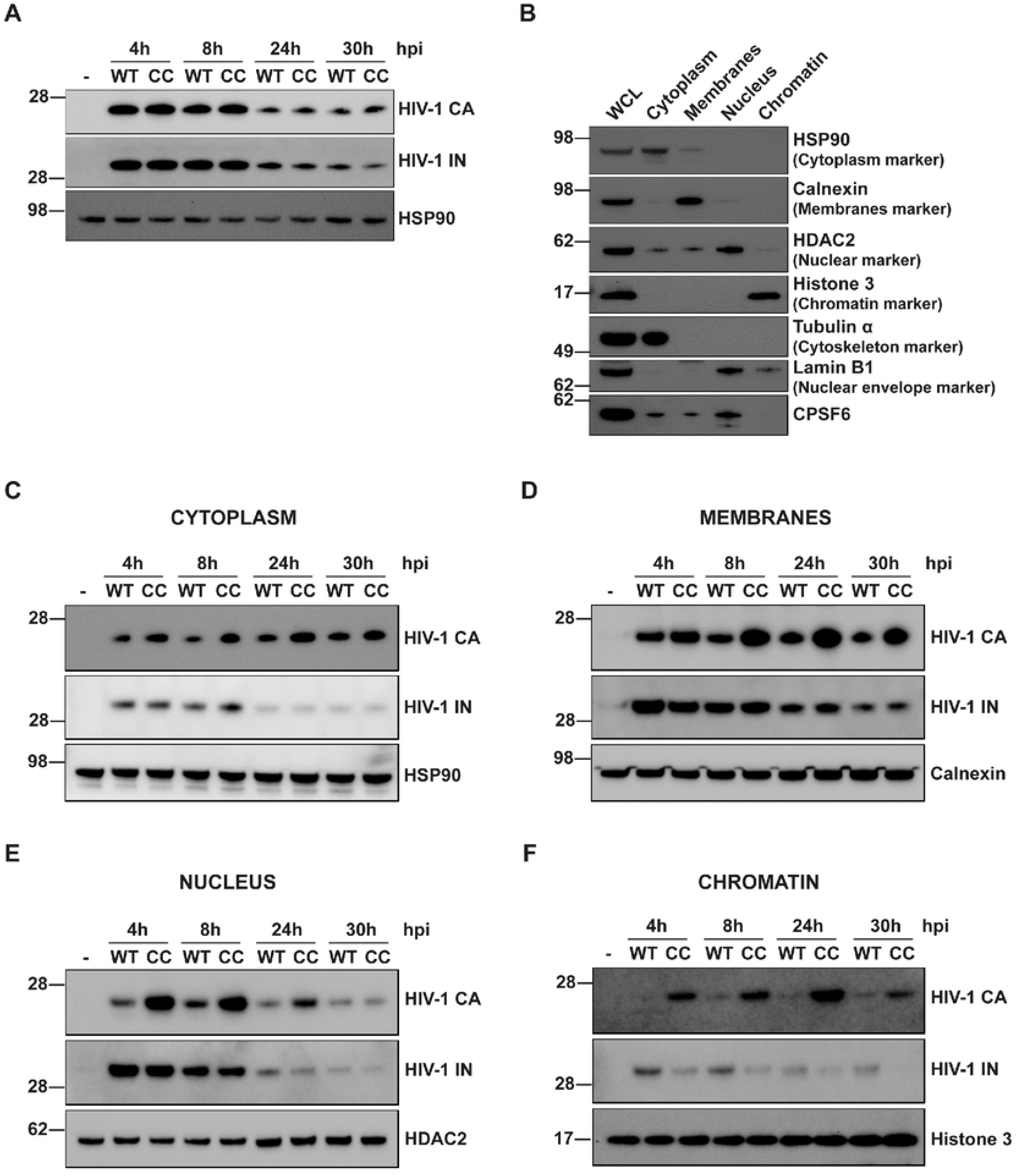
Cellular localisation of CA and IN proteins during WT and A14C/E45C infections. HeLa cells were synchronously infected with equal RT units of WT or A14C/E45C mutant (CC in the figure) VLP. At 4, 8, 24 and 30 h post-infection, cells were harvested in parallel to be processed as a whole cell lysate or to undergo subcellular fractionation. Protein levels were quantified by BCA assay, proportional amounts of the fractions related to the WCL were loaded on SDS-PAGE gels and analysed by immunoblotting using the following antibodies: Anti-CA and anti-IN for HIV-1 proteins, HSP90 for cytoplasm, Calnexin for membranes, HDAC2 for nucleus, Histone 3 for chromatin and Tubulin α for cytoskeleton. Lamin B1 was used as a nuclear envelope marker. “-“ indicates uninfected cells. Panels show representative immunoblots of: (A) WCL probed for HIV-1 CA and IN. HSP90 was blotted as a loading control. (B) Subcellular fractions from uninfected cells, probed for control proteins to confirm successful sample fractionation. (C-E) Subcellular fractions probed for HIV-1 CA and IN and the appropriate cellular marker: (C) cytoplasm, (D) membranes, (E) nucleus and (F) chromatin fraction.

### The A14C/E45C mutant is impeded at nuclear entry

In order to investigate nuclear entry of the A14C/E45C mutant further, we monitored the interactions between CA and the nuclear pore proteins Nup358 and Nup153 during infection. These proteins are part of the nuclear pore complex (NPC) where Nup358 faces the cytoplasm and Nup153 forms part of the nuclear basket of the NPC, facing the nucleus. Furthermore, both proteins have previously been described to bind HIV-1 CA directly and to be involved in HIV-1 replication [25, 26, 30]. HeLa cells were synchronously infected with equal RT units of WT or A14C/E45C VLP for 2, 4, 6, 8 and 10 hpi. At each time point, cells were fixed, and proximity ligation assays (PLA) were performed with specific antibody combinations: anti-HIV-1 CA and anti-Nup358 (Fig 7A-C), and anti-HIV-1 CA and anti-Nup153 (Fig 7D-F). In this assay, direct protein-protein interactions or proteins spaced < 40 nm apart can be visualised as foci by immunofluorescence. In addition, by doing *post hoc* analysis, the localisation of these interactions can be analysed in relation to the nuclear envelope, in this case approximated by DAPI staining. Fig 7A shows representative cells assayed for CA-Nup358 co-localisation at 8 hpi. Counting the number of foci per cell (Fig 7B) revealed that there were similar levels of interaction between CA and Nup358 throughout the time course of infection, and there was no significant difference between WT and A14C/E45C infections. The localisation of these foci (Fig 7C) seemed to be around the DAPI edge for both VLP, but with a tendency for WT foci to be further into the nucleus than A14C/E45C foci. This suggests that there is little difference in the early stages of infection, up to reaching the nuclear pore, between WT and A14C/E45C VLP, and, once again, that cores reach the nucleus within 2 hpi, sooner than previously expected. In contrast, there were clear differences in the staining for CA-Nup153 co-localisation between WT and A14C/E45C infections (Fig 7D-F). Analysis of the number of foci per cell showed that there was increased A14C/E45C foci compared to WT infections at all time points (Fig 7E). As equal numbers of cores were arriving at the nucleus, as measured by CA-Nup358 staining (Fig 7B), this suggests that the A14C/E45C cores were spending more time at the nuclear pore. Moreover, the A14C/E45C foci appeared to localise at the cytoplasmic side of the DAPI edge while the WT foci were mainly on the nuclear side (Fig 7F). Given that a layer of lamin and the nuclear envelope surround the chromatin in the nucleus, the DAPI edge does not indicate the exact nuclear/cytoplasmic boundary, but rather the inner side of the nuclear pore. Thus, being on the cytoplasmic side of the DAPI staining, together with increased CA-Nup153 staining, suggests that A14C/E45C hyper-stable cores are being trapped at the NPC.

**Figure 7.**
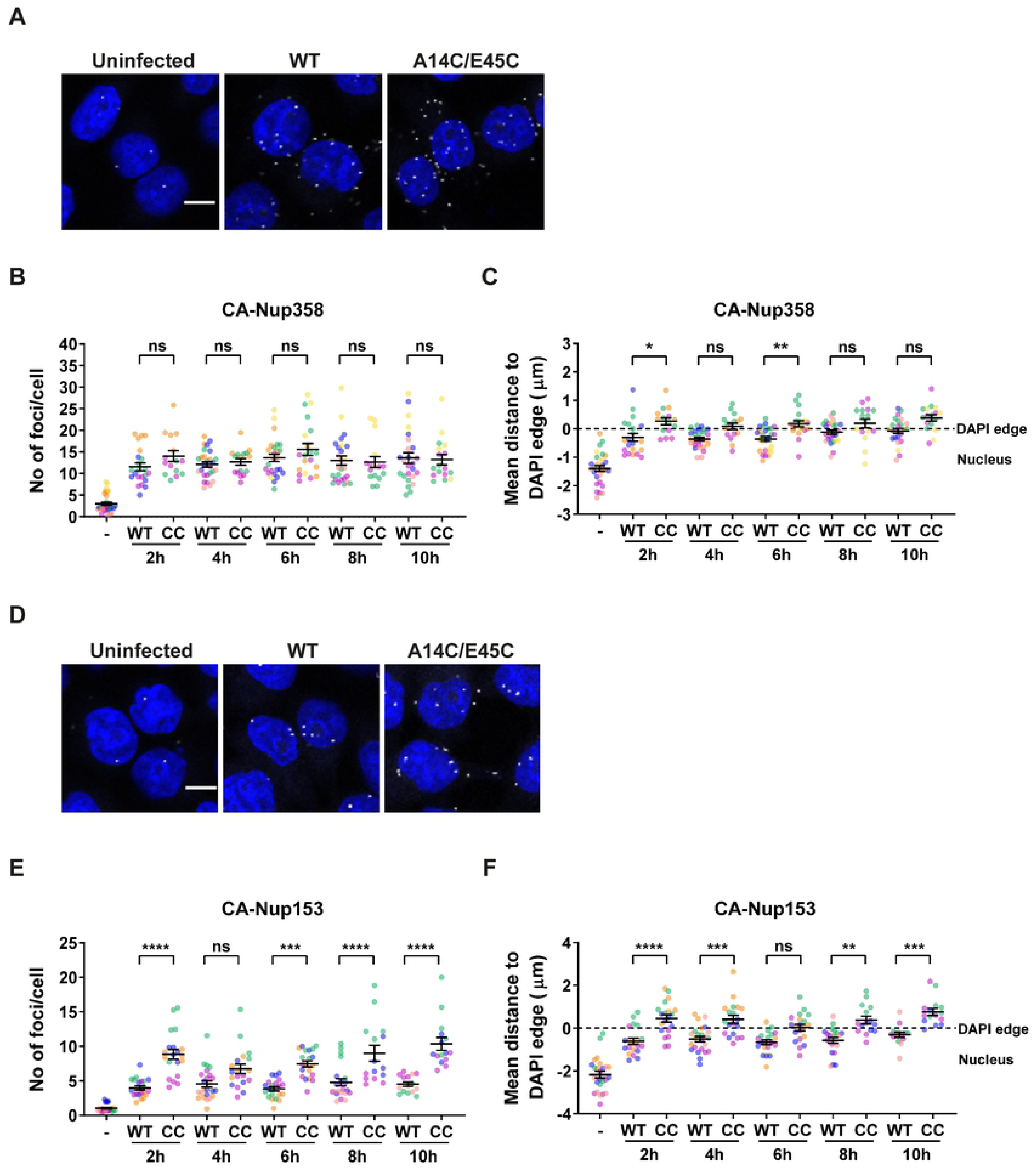
CA co-localisation of CA with Nup358 and Nup153 during infection. HeLa cells were synchronously infected with equal RT units of WT or A14C/E45C VLP and fixed at 2, 4, 6, 8 and 10 hpi. Cells were incubated with primary antibodies to HIV-1 CA and either Nup358 (A-C) or Nup153 (D-F), followed by specific secondary antibodies conjugated to the PLUS and MINUS PLA oligonucleotides (probes). Each foci represents a positive PLA signal generated by the amplication of the interaction between the PLUS and MINUS probes. (A) and (D) show representative images of CA-Nup358 and CA-Nup153 co-localisation at 8 hpi, respectively (the scale bar is 10µm). Longitudinal Z-series were acquired with a 63X objective using a confocal microscope followed by 3D image analysis performed with the GIANI plug-in in FIJI. The number of foci per cell (B and E) and the distance of those foci to the DAPI edge (C and F) were quantified. In the plots, each point represents the mean foci data from all the cells within an image (75 to 100 cells/image). At least 12 images were acquired per condition over at least three independent biological repeats (plotted in different colours). Overall means ± SEM are shown in black. *p<0.05, **p<0.01, ***p<0.001, ****p<0.0001, ns = not significant.

### CPSF6 is not re-localised to nuclear speckles during infection with hyper-stable mutants

In addition to binding nuclear pore proteins, CA interacts with CPSF6, which has been reported to direct HIV-1 integration site specificity. Specifically, CPSF6 is proposed to locate HIV-1 to highly transcribing regions of the genome identified as nuclear speckles, or speckle-associated domains (SPADs) [9, 34, 35]. The absence of CPSF6, or blocking of CPSF6-CA binding, promotes integration into lamina-associated domains (LADs), regions of heterochromatin near the nuclear envelope [9, 34]. Unfortunately, despite repeated attempts, we were unable to optimise our PLA assay to reliably visualise CA-CPSF6 co-localisation. However, recently, it has been observed that CPSF6 only redistributes to SPADs during WT HIV-1 infection and not during infection with CPSF6-binding deficient mutants A77V and N74D [35, 48]. This suggests that the CA-CPSF6 interaction is driving CPSF6 reorganisation in the nucleus. Therefore, to investigate the interaction between our hyper-stable mutants and CPSF6, we monitored the reorganisation and redistribution of CPSF6 to SPADs during infection. HeLa cells were synchronously infected with equal RT units of WT, A14C/E45C, E180C or M68C/E212C VLP. At 16 hpi, cells were fixed and immuno-stained with antibodies against HIV-1 CA, CPSF6 and SC35, also called serine and arginine rich splicing factor 2 (SRSF2), which is a marker for SPADs [62]. Due to the species of the antibodies, we could only co-stain for pairs of markers at a time. Fig 8A and Fig S3 show that cells infected with WT-HIV-1 exhibit a compelling redistribution of CPSF6 into puncta that co-localise with SC35-positive nuclear speckles, confirming previous reports [35, 48]. However, there was no such redistribution of CPSF6 during infection with any of the hyper-stable mutants which showed similar CPSF6 staining to uninfected cells. Importantly, when we co-stained for CA and CPSF6, ∼80% of WT CA positive cells contained CPSF6 puncta (Fig 8B), compared to only 5% of hyper-stable CA positive cells. Although we were able to detect some CA signal in the nucleus of WT HIV-1 infected cells that colocalised with the CPSF6 signal by immunofluorescence (Fig 8B), the CA signal was relatively weak and we cannot say whether this CA was part of a core or not. The CA staining in the nucleus was not as evident during infection with the hyper-stable mutants. Together, these data show that WT CA is able to interact with CPSF6 and induce its redistribution to SPADs, whilst the hyper-stable CA fail to alter CPSF6 localisation.

**Figure 8.**
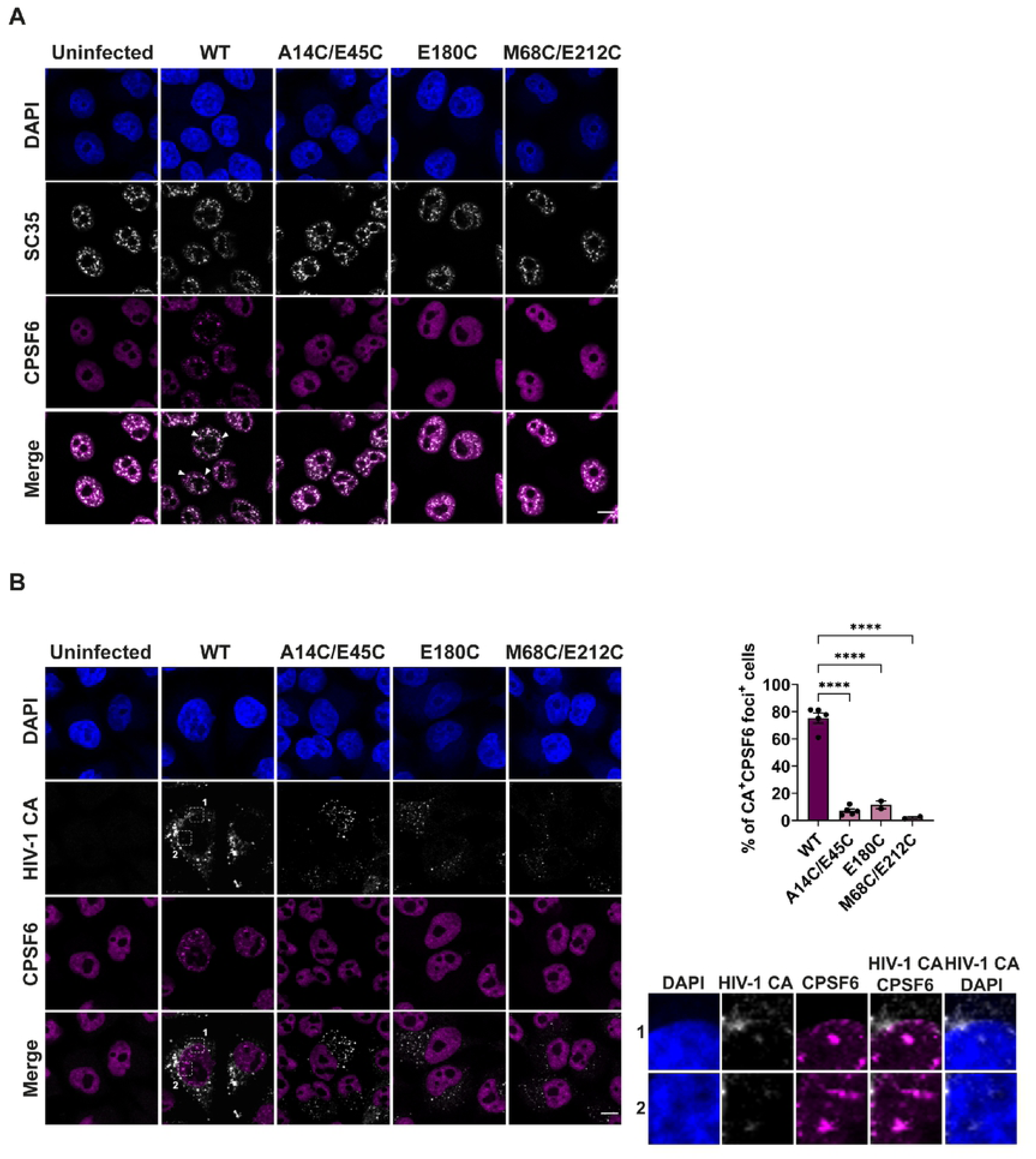
HIV-1 CA, CPSF6 and SC35 staining during infection with WT and hyper-stable mutant VLP. HeLa cells were synchronously infected with equal RT units of WT, A14C/E45C, E180C or M68C/E212C VLP and fixed at 16hpi. Cells were incubated with primary antibodies against HIV-1 CA, CPSF6 and SC35 followed by specific secondary antibodies conjugated to Alexa Fluor fluorophores. (A) Representative images of SC35 and CPSF6 staining. White arrows point to SC35 and CPSF6 co-localisation in the merged WT image. (B) Representative images of HIV-1 CA and CPSF6 staining (left panels). White boxes in WT columns indicate zoomed-in regions labelled 1 and 2 (right panels). The scale bar in (A) and (B) represents 10µm. Bar chart shows the number of CA positive cells that showed CPSF6 redistribution to puncta. Points indicate individual biological repeats (∼150 cells per experiment were used for quantification) and lines show the mean ± SEM; ****p<0.0001.

## DISCUSSION

It has been known for many years that HIV-1 can infect non-dividing cells and so must cross the nuclear envelope during infection. Until recently, the dogma was that HIV-1 cores uncoated in the cytoplasm during reverse transcription, because they were too large to cross the nuclear pore. However, new evidence has led people to re-examine this order of events. When and where the CA lattice breaks down is currently highly controversial. Theories range from cytoplasmic uncoating to uncoating at the nuclear pore (reviewed in [63]) [35, 38, 39, 42, 64], and more recently, uncoating inside the nucleus [43, 44, 48, 65]. Along these lines, emerging electron microscopy images appear to show intact cores going through nuclear pores [65]. Some open questions are (1) Can reverse transcription complete inside a core? (2) Can a complete core cross the nuclear pore? (3) What replication events require CA? and (4) When is uncoating required? Unfortunately, uncoating is difficult to measure directly. Therefore, in this study, we have taken a genetic approach to investigating uncoating by comparing the replication of WT HIV-1 with hyper-stable CA mutants that are potentially slower or unable to uncoat. Thus, we have not asked “where/when does uncoating occur?”, but rather “where/when does viral replication get blocked if uncoating is prevented?”.

We compared a panel of mutants designed to induce disulphide bonds at different CA lattice interfaces. Many of these have previously been used for structural studies and clearly induced stability of *in vitro* assembled CA [7, 8, 11, 12, 57, 66–69], but anecdotally were thought unlikely to make stabilising disulphide bonds in the reducing environment of the cytoplasm. Nevertheless, we found that two previously described intra-hexamer mutants, A14C/E45C and M68C/E212C [11, 12, 57], and a novel inter-hexamer mutant, E180C, all showed increased lattice stability in cells (Fig 2A and Fig 6) and maintained disulphide bonds following infection (Fig 5A). Interestingly, these three mutants represent stabilising different interfaces within the CA lattice. Thus, it appears that no particular interface is dominant in lattice stability. It was not surprising that most of the mutants did not increase core stability, as there is a lot of variation in the exact Cβ-Cβ distances between individual residues, depending on the region of the lattice that they reside (i.e. depending on the curvature of that part of the fullerene cone [5, 66]). This is especially true around the trimeric interface. It is intriguing that the NTD-NTD mutant A14C/E45C resulted in hyper-stable cores whilst the NTD-NTD mutant A42C/T54C did not, as they have near identical hexamer crystal structures [12, 67]. However, the geometry and exact chemistry of the surrounding environment likely affect whether disulphide bonds form in cells.

### Uncoating is required between reverse transcription and integration

As expected, although they produced normal viral titres as measured by particle release (Fig S1), most of the mutants in the panel had reduced infectivity (Fig 3). The intra-hexamer mutants were the least infectious, perhaps highlighting the importance of residues within the CA N-terminal domain for early replication steps. The severity of the infectivity defect did not correlate with core stability, as measured by the fate-of-CA assay, but did generally reflect the level of reverse transcription (Fig 4G and H). The exceptions to this were the three mutants identified as hyper-stable. Surprisingly, despite uncoating previously being linked to reverse transcription [38, 49, 51, 61, 70], A14C/E45C and E180C were able to reverse transcribe to approximately WT levels and, furthermore, showed only a partial reduction in 2-LTR circle production (Fig 4 and Fig 5B). This shows that reverse transcription can complete in a hyper-stable core and implies that it is not necessary for the core to uncoat in order to accommodate the double stranded DNA. These results are in line with previous studies showing that hyper-stable mutants E45A, Q63A/Q67A, 5Mut (Q67H/K70R/H87P/T107N/L111) and A14C/E45C can reverse transcribe [48, 52, 71, 72] and with a recent report saying that reverse transcription can complete in whole WT cores [43]. However, it is possible that the hyper-stable cores are partially opened. Indeed, the production of 2-LTR circles would imply that the core was open to some extent in order for the cellular ligases to access the viral DNA. Recent *in vitro* studies have suggested that reverse transcription increases the pressure inside the core [61], inducing mechanical changes in the capsid that progressively remodel the lattice [73, 74] and may result in viral DNA loops bursting out of partially uncoated cores [75]. This agrees with our previous assessment that uncoating is triggered after first strand transfer during reverse transcription [51]. By inducing disulphide bond formation, we may have prevented some or all of these remodelling changes occurring in the hyper-stable mutant cores, and therefore prevented more extensive uncoating from occurring. This implies that whilst reverse transcription promotes uncoating, it is not dependent on the disassembly of the lattice. Instead, the virus appears to be using reverse transcription to time uncoating.

Nonetheless, hyper-stability can be detrimental for reverse transcription as the M68C/E212C mutant showed a 10-fold decrease in reverse transcription products compared to WT (Fig 4D). The cysteine residues introduced in M68C/E212C are located on the intra-hexamer NTD-CTD interface, which is important for the curvature of the lattice. Christensen *et al*. recently reported that the compound GS-CA1, which binds at the NTD-CTD interface, affected capsid integrity and strongly inhibited reverse transcription *in vitro*, and proposed that affecting the NTD-CTD interface introduces lattice strain and promotes capsid fracturing [75]. Thus, it could be possible that the formation of disulphide bonds at this specific interface affects the structure of the CA lattice in such a way that is not compatible with reverse transcription.

Importantly, even the mutants that reverse transcribed well were still markedly impaired for integration (Fig 5B), presumably meaning that the cores are unable to open fully, or in the correct manner to allow integration. This agrees with various recent reports suggesting that a final capsid uncoating reaction needs to occur before integration can take place [35, 39, 42–44, 48, 64, 65]. Stabilising the CA lattice by either mutation, or drugs, probably affects flexibility as well as lattice break down, so it is hard to separate whether the core needs to restructure in some way or to break apart completely. We therefore use the term “CA remodelling” to cover both of these events.

The link between reverse transcription and uncoating has recently taken a new twist, as the location of reverse transcription in the cell has been questioned. There is increasing evidence that CA-containing viral complexes reach the nucleus early, before reverse transcription has completed [35, 43, 44, 48, 76, 77], and that reverse transcription actually completes in the nucleus [44]. In support of nuclear-associated reverse transcription, here, we also detected CA and, importantly, IN in nuclear fractions at 4 hpi (Fig 6E) and detected CA interacting with nuclear pore proteins at 2 hpi (Fig 7), well before the peak of reverse transcription at 6hpi (Fig 4).

### Hyper-stable CA mutants are compromised for nuclear entry

As two of our hyper-stable mutants seemed to have a major defect after reverse transcription, we asked whether these hyper-stable cores could enter the nucleus. 2-LTR circles are considered a surrogate for nuclear entry [78, 79], and as A14C/E45C and E180C produced 2-LTR circles, albeit with a 5-fold decrease compared to WT (Fig 5B), it suggested that they could indeed enter the nucleus. Our subcellular fractionation experiments confirmed that both WT and hyper-stable CA were detected in all fractions following infection, including both soluble and chromatin-associated nuclear fractions (Fig 6). To confirm where the nuclear envelope fractionated, we blotted for lamin B1 which is located inside the inner layer of the nuclear envelope. Lamin B1 was detected in the nucleus and, to a lesser extent, in the chromatin fraction (Fig 6B), suggesting that the CA in these fractions could be at the nuclear envelope as well as inside the nucleus. Importantly, chromatin is known to be tightly linked with nuclear pores and lamin [80]. There was noticeably less WT HIV-1 CA present at each time point than the hyper-stable mutant CA, particularly in the nuclear fractions, suggesting that it was turned over faster (Fig 6E and F), although it is also possible that WT CA was more susceptible to degradation during the fractionation process. Conversely, the levels of IN were similar at all time points between WT and hyper-stable infections in all fractions except the chromatin associated fraction, where there was more IN detected following WT infections. As WT CA was barely detectable here (Fig 6F), this could reflect WT cores being able to breakdown and deliver IN more efficiently to the chromatin than A14C/E45C cores, and that once disassembled, the CA is degraded. Together, these data show that although the hyper-stable cores retain more CA than WT cores, the IN levels are comparable until the cores meet chromatin. This implies that uncoating has little impact on replication until a chromatin associated event.

To explore whether hyper-stable cores could truly enter the nucleus or were associating with nuclear membranes, we studied the dynamics of WT and A14C/E45C CA interactions with nuclear pore proteins (Fig 7). We chose Nup358 and Nup153 because they are located on the outer and inner face of the nuclear pore, respectively, and because their interaction with CA is well characterised [25, 26, 29, 30]. We found that WT and A14C/E45C CA showed similar levels of interaction with Nup358 over time (Fig 7B) suggesting that both cores could reach the nuclear pore with equivalent kinetics, agreeing with our fractionation experiments. As seen in a previous report [81], we observed some CA-Nup358 foci in the cytoplasm, which was more noticeable during A14C/E45C infection, but there were no significant differences between the WT and mutant CA distribution (Fig 7C). In contrast, there was clear divergence in the co-localisation of CA and Nup153 between WT and hyper-stable cores. There were significantly increased numbers of foci for the A14C/E45C CA at all time points. As equal numbers of cores were arriving at the nucleus (as measured by Nup358 co-localisation), this suggests that the A14C/E45C cores were spending more time at the nuclear pore. Moreover, the A14C/E45C foci appeared to localise more towards the cytoplasmic side of the DAPI edge while the WT foci were mainly on the nuclear side, suggesting that WT cores travel further into the nucleus and that A14C/E45C hyper-stable cores are trapped at the NPC. It is worth pointing out that quantifying the distance of the PLA foci to the DAPI edge has caveats. The PLA signal is the result of a complex of rolling circle amplification products together with antibodies bound to two proteins of interest. As this is a big complex, the foci distance to the DAPI edge cannot be taken as an absolute distance. However, the same caveats apply to both WT and mutant cores, so a relative comparison can be made. Thus, it seems that uncoating or lattice flexibility is required for nuclear entry and points to a CA remodelling event at the nuclear pore, as suggested by some other labs [42, 65]. It is interesting to recall that the A14C/E45C mutant still produces 2-LTR circles, despite apparent limited nuclear entry, questioning what this product really represents.

Interestingly, studies with the CPSF6 binding-defective HIV-1 CA mutants, N74D and A77V, have reported that a longer residence time at the nuclear envelope promotes integration into heterochromatin regions close to the nuclear envelope [9, 34, 35, 43, 48]. Furthermore, unlike WT HIV-1, neither the N74D nor the A77V mutant can re-localise CPSF6 to SPADs, showing that this is a CA dependent event [9, 34, 35, 48]. Since A14C/E45C cores were stalled at the nuclear pore, we were intrigued to see whether this mutant would be able to promote CPSF6 redistribution to nuclear speckles. In agreement with very recent reports [35, 48], we found that only WT-infected cells showed a redistribution of CPSF6 to SC35 positive puncta (Fig 8). Importantly, A14C/E45C is still able to bind CPSF6 *in vitro* [82, 83], suggesting that either it does not have access to CPSF6 in cells, or that it is unable to relocate to nuclear speckles because it is retained elsewhere. CPSF6 binds to the same pocket on the CA lattice as Nup153 [29, 31, 32] and Bejarano *et al* have recently reported the consecutive binding of the hexameric CA lattice to Nup153 and then CPSF6 in macrophages [40]. Thus, we speculate that if A14C/E45C is still binding Nup153 at the nuclear pore, it might not be able to uncouple and move on to binding CPSF6. We hypothesize that the CA lattice needs to remodel at the pore in order to be released from Nup153 and move into the nucleus, where it can then interact with CPSF6 and move to an optimal site for integration. As CA is required for the CPSF6 interaction, it follows that a fraction of CA must be retained by the pre-integration complex until chromatin binding, but sufficient CA must be removed to allow IN and the viral cDNA to access chromatin for integration itself to occur.

### Models of HIV-1 uncoating

We have illustrated the current models of HIV-1 uncoating in Fig 9. From left to right, uncoating in the cytoplasm, uncoating at the nuclear pore, uncoating inside the nucleus or uncoating at the integration site. Early uncoating or a failure to uncoat both have detrimental effects on virus infectivity. The data presented here point to a remodelling event at the nuclear envelope that, although it is not required for reverse transcription to complete, is needed to completely pass through the nuclear pore, allow binding to CPSF6 and, ultimately, for integration. We therefore think it likely that some sort of uncoating event begins at the nuclear pore and finishes at the site of integration. We have also previously shown that the murine leukaemia virus (MLV) p12 protein binds directly to both CA and nucleosomes, tethering the MLV core to chromatin during mitosis, and have proposed that p12 could be acting in a similar capacity to CPSF6 in HIV-1 infection [84]. Figure 9 includes our model of MLV uncoating. As MLV cannot traverse nuclear pores [85] it is unlikely that it undergoes a similar pore-induced CA remodelling event, suggesting that MLV uncoating might be triggered by chromatin binding. As MLV must wait for mitosis, it is possible that an intact CA lattice is needed until integration to provide a protective environment for the viral cDNA.

**Figure 9.**
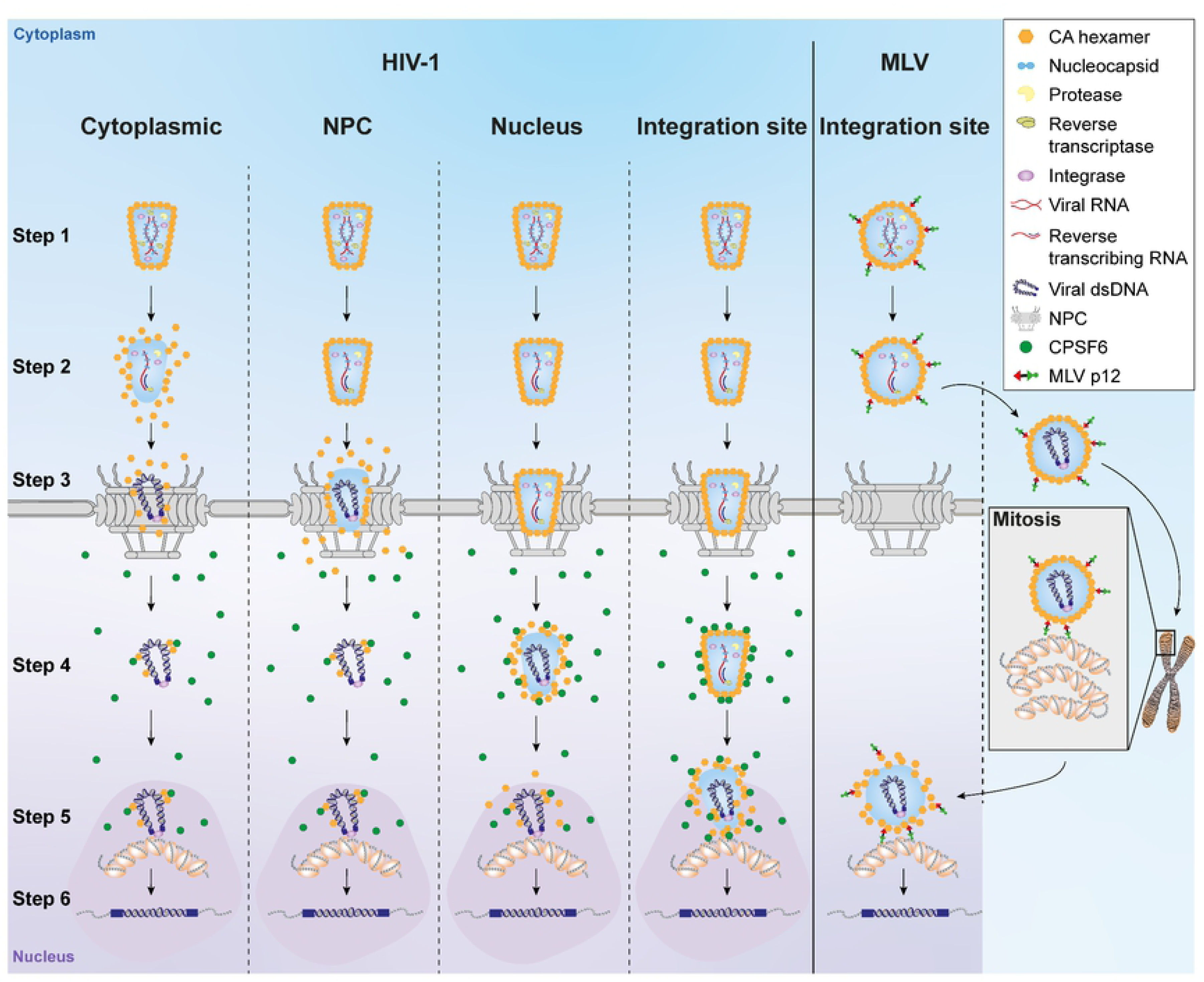
Current models for HIV-1 and MLV uncoating. Incoming viral cores (HIV-1, left columns and MLV, right column) surrounded by a CA lattice (orange hexamers) contain the viral RNA (red) coated with nucleocapsid (light blue), as well as the viral enzymes, protease (yellow), reverse transcriptase (light green) and integrase (purple) (step 1). Reverse transcription of the RNA to double stranded DNA (dark blue) starts following infection and the core travels towards the nucleus down microtubules (step 2). HIV-1 DNA can cross the nuclear pore (step 3), and in association with CPSF6 (green dots) (step 4), move to nuclear speckles (darker pink shaded regions) and integrate into the host cell chromatin (grey with orange ovals) (step 5) to form a provirus (step 6). When and where the CA lattice disassembles and where reverse transcription finishes is still debated and the possible scenarios for HIV-1 uncoating (loss of orange hexagons) are illustrated here: from left to right; uncoating in the cytoplasm, uncoating at the nuclear pore, uncoating inside the nucleus or uncoating at the site of integration. In contrast, MLV cannot pass through nuclear pores and must wait for mitosis to occur before accessing the chromatin. Our current model for MLV uncoating is shown on the right: At mitosis, the N-terminal domain of the MLV p12 protein (red) binds directly to CA and the C-terminal domain (green) binds to nucleosomes, tethering the likely intact MLV core to the host chromatin. Following mitosis, MLV uncoats to allow integration, which, as it is already associated with chromatin, probably occurs at the integration site.

In conclusion, we have demonstrated that HIV-1 with a hyper-stable CA lattice is able to reverse transcribe successfully but is stalled at nuclear entry, which has a negative effect on CPSF6 binding and integration. We suggest that an uncoating or CA remodelling event normally occurs at the nuclear pore and that this is essential for the later replication events that take place in the nucleus. Furthermore, our observations suggest that viral cores are present at the nucleus before reverse transcription is completed. Therefore, it is plausible that both reverse transcription and uncoating finish in the nucleus. Further work is needed to fully understand the state of the CA lattice in the nucleus and the exact relationship between uncoating and reverse transcription.

## MATERIAL & METHODS

### Cell lines

Adherent cell lines, 293T, HeLa and Vero cells, were maintained in Dulbecco’s modified Eagle medium (Thermo Fisher), and suspension cell lines, SupT1 and U937 cells, were maintained in RPMI-1640 (Thermo Fisher). All cell lines were authenticated and tested mycoplasma-free from Bishop laboratory cell stocks. Media was supplemented with 10% heat-inactivated foetal bovine serum (FBS; Biosera) and 1% Penicillin/Streptomycin (Sigma). Cells were grown in a humidified incubator at 37°C and 5% CO_2_.

### Plasmids and site-directed mutagenesis

The plasmids used to produce HIV-1 VLP, pVSV-G, pCMVΔR8.91 and pCSGW, have been described previously [86, 87]. To create Gag-Pol plasmids carrying cysteine-substitution mutations in CA, site-directed mutagenesis was performed on pCMVΔR8.91 using the QuickChange II-XL site-directed mutagenesis kit (Agilent) according to manufacturer’s instructions and using the primers listed in Supplementary Table 1. Repeated site-directed mutagenesis was performed to create the double, triple and quadruple mutants. The introduction of the desired mutations was confirmed by Sanger sequencing (Source Bioscience).

### Virus-like particle (VLP) production

HIV-1 virus-like particles (VLP) were produced by co-transfecting 293T cells with a 1:1:1 ratio of three plasmids: pVSV-G, pCSGW (GFP-reporter) and pCMVΔ8.91 (or pCMVΔ8.91 mutants). Approximately 16h post-transfection, cells were treated with 10mM sodium butyrate for 8h and VLP-containing supernatants were harvested 24 h later. VLP titres were analysed using a Lenti RT ELISA kit (Cavidi) following manufacturer’s instructions. For the fate-of-capsid, cell fractionation and PLA assays, VLPs were concentrated by ultracentrifugation through a 20% (w/w) sucrose cushion using a Beckman SW32Ti rotor (Beckman-Coulter) at 20,200 rpm for 2h at 4°C.

### Single round infectivity assay

293T, HeLa, SupT1 and U937 cells were challenged with normalised amounts of WT and mutant VLP based on their RT activity and incubated for 72 h at 37°C. The percentage of GFP-expressing cells was analysed by flow cytometry using a FACS VERSE, LSR or Fortessa A analysers (BD Biosciences). Data was analysed using FlowJo software.

### Immunoblotting

Cells were lysed in ice-cold radioimmunoprecipitation assay (RIPA) buffer (Thermo Fisher) supplemented with a protease inhibitor cocktail (Roche) and proteins were separated on 4-12% or 10% Bis-Tris SDS-PAGE gels (Thermo Fisher). To assess disulphide cross-linking in cells, 4x SDS sample buffer (Thermo Scientific) was added to cell lysates prior to treatment with 50mM Iodoacetamide (Sigma) or 0.1M DTT (Sigma). Samples treated with Iodoacetamide were incubated for 30min at room temperature followed by a 15 min incubation at 65°C. Samples treated with DTT were boiled at 95°C for 5 min. Samples were then applied to 8% Bolt Bis-Tris Plus gels (Thermo Fisher) and electrophoresed in Bolt MOPS buffer (Thermo Fisher). Primary antibodies used were: anti-HIV-1 CA (in house), anti-HIV-1 IN (in house), anti-HSP90 (CST; #4874), anti-Calnexin (CST; #2679), anti-HDAC2 (CST; #5113), anti-Histone 3 (CST; #14269), anti-Tubulin-α (Bio-Rad; VMA00051) and anti-Lamin B1 (Proteintech; 66095-1-Ig). Secondary antibodies used were: anti-mouse and anti-rabbit HRP-conjugated secondary antibodies (Thermo Fisher; 61-6520 and 31460); anti-mouse IRDye 800CW secondary antibodies (LICOR). Blots were analysed on a Chemidoc imaging system (Bio-Rad) or by X-ray film exposure or on an Odyssey CLx imaging system (LICOR).

### Fate-of-capsid assay

The fate-of-capsid assay was performed as previously described (Yang *et al*, 2014) but with some modifications. HeLa cells were seeded at 10^6^ cells/well in 6-well plates one day prior infection and spinoculated (1600rpm at 16°C for 30min, followed by 37°C for 1h) with equal amounts of VLP based on their RT activity. Cells were harvested at 2 and 20 hpi and cell pellets were lysed by adding hypotonic buffer (10mM Tris-HCl pH 8.0, 10mM KCl and 1mM EDTA pH 8.0 supplemented with a protease inhibitor cocktail (Roche)) and passing through a Qiashredder column (Qiagen). At this point, 50 µL was harvested as the input (I) and resuspended in 2x Laemmli’s SDS-PAGE sample loading buffer (Sigma). The remaining lysate was layered on top of a 30% (w/w) sucrose cushion and centrifuged using a Beckman SW41 rotor (Beckman Coulter) at 32,000rpm and 4°C for 1 h, to separate soluble and assembled CA. After centrifugation, 500 µL of the uppermost portion of the supernatant was harvested as the soluble fraction (S). Following aspiration of the sucrose cushion, the pellet (P) was resuspended in 100 µL of 2x sample loading buffer. The soluble fraction was precipitated using methanol-chloroform extraction and resuspended in 100 µL of 2x sample loading buffer. Then, the I, S and P samples were analysed by western blotting with an anti-HIV-1 CA antibody.

### Quantitative PCR analysis to measure RT products

Quantitative PCR analysis was conducted as previously described (Bishop *et al*, 2008; Cosnefroy *et al*, 2016). Briefly, VLP were treated with 20 units/mL RQ1-DNase (Promega) in 10mM MgCl_2_ for 1h at 37°C before infection. 293T cells were spinoculated (1600rpm at 16°C for 30min) followed by a 30min incubation at 37°C. The reverse transcriptase inhibitor Nevirapine (NVP; Sigma) was used as a negative control, prepared in DMSO and used at a final concentration of 10 µM. Cells were harvested at the indicated times post-infection and total DNA was extracted using the DNeasy Blood & Tissue kit (Qiagen). The extracted DNA was digested with 1 unit/µL DpnI (Thermo Fisher) for 2.5 h at 37°C. qPCR was performed in TaqMan real-time PCR master mix (Thermo Fisher) with 900 nM primers and 250 nM probes. The reactions were performed on a 7500 fast real-time PCR system (Applied Biosystems). To calculate DNA copy numbers, standard curves were generated from serial dilutions of pCSGW or p2-LTR junction in 293T cellular DNA. The following primers and probes were used; strong stop cDNA products: *for* 5’-*TAACTAGGGAACCCACTGC*, *rev* 5’-*GCTAGAGATTTTCCACACTG* and *probe* 5’-*FAM-ACACAACAGACGGGCACACACTA-TAMRA*; second strand cDNA products: *for* 5’-*TAACTAGGGAACCCACTGC*, *rev* 5’-*CTGCGTCGAGAGAGCTCCTCTGGTT* and *probe* 5’-*FAM-ACACAACAGACGGGCACACACTA-TAMRA*; 2-LTR junction: *for* 5’-*GTGTGTGCCCGTCTGTTG*, *rev* 5’-*CAGTACAAGCAAAAAGCAGATC* and *probe* 5’-*FAMGGTAACTAGAGATCCCTCAGACCTAMRA*.

### Cell fractionation assay

HeLa cells were infected with equal amounts of WT and mutant VLP (based on their RT activity) by spinoculation (1600rpm, 16°C for 2h) followed by a 30 min incubation at 37°C. At the indicated times post-infection, cells were harvested in parallel either as a whole cell lysate (WCL) or for processing with the subcellular protein fractionation kit for cultured cells (Thermo Fisher) following manufacturer’s instructions. Protein content in the WCL and in the different fractions was measured by BCA assay (Thermo Fisher) using a FLUOstar Omega plate reader (BMG Labtech). Relative amounts of each fraction compared to the WCL were analysed by immunoblotting.

### Immunofluorescence

HeLa cells were seeded at 0.8x10^5^ cells/well on 13 mm glass coverslips in 24-well plates the day prior to infection. Cells were infected with equal RT units of WT or mutant VLP by spinoculation (1600rpm, 16°C for 2h) followed by a 30min incubation at 37°C. At the indicated times post-infection, cells were washed twice with ice-cold PBS, fixed with PBS supplemented with 4% paraformaldehyde for 5min at room temperature followed by an ice-cold methanol incubation for 5min at −20°C, and washed twice again with ice-cold PBS. After permeabilization with 0.5% saponin (Sigma) in PBS for 30min at room temperature, cells were blocked in 5% donkey serum (DS; Sigma) and 0.5% saponin in PBS for, at least, 1h at room temperature. Then, cells were incubated with the following primary antibodies: anti-HIV-1 CA (in house), anti-CPSF6 (Atlas; HPA039973) and anti-SC35 (Abcam; ab11826), diluted in 1% DS with 0.5% saponin in PBS (antibody buffer) for 1h at RT. After three washes with PBS, cells were incubated with the following secondary antibodies: goat anti-mouse-AF488 (abcam; ab150117), donkey anti-rabbit-AF568 (abcam; ab175692), donkey anti-mouse-AF647 (Thermo Fisher; A-31571) and donkey anti-rabbit-AF647 (Thermo Fisher; A-31573) in antibody buffer for 1h at RT. After three washes with PBS, the coverslips were mounted on glass slides with ProLong gold antifade mountant with DAPI (Thermo Fisher).

### Proximity ligation assay (PLA)

HeLa cells were seeded at 10^5^ cells/well on 13mm glass coverslips in 24-well plates the day prior to infection. Cells were infected with equal RT units of WT or mutant VLP by spinoculation (1600rpm, 16°C for 2h) followed by a 30min incubation at 37°C. At the indicated times post-infection, cells were washed twice with ice-cold PBS, fixed with PBS supplemented with 4% paraformaldehyde for 5min at room temperature followed by an ice-cold methanol incubation for 5min at −20°C, and washed twice again with ice-cold PBS. Then the immunofluorescence protocol was followed using the following pairs of primary antibodies: anti-HIV-1 CA (in house) and anti-Nup358 (Abcam; ab64276); anti-HIV-1 CA (in house) and anti-Nup153 (Abcam; ab84872), diluted in 1% DS with 0.5% saponin in PBS for 1h at RT. From this point, the Duolink PLA fluorescence detection kit’s protocol (Sigma) was followed. After three washes with PBS, coverslips were incubated with secondary antibodies conjugated to PLA probes (anti-mouse PLUS or anti-rabbit MINUS; Sigma) in antibody buffer (Sigma) for 1h at 37°C. Coverslips were incubated with ligase for 30 min at 37°C followed by amplification by polymerase for 100 min at 37°C. From the primary antibody incubation, coverslips were washed with buffers A and B (Sigma) as indicated in the manufacturer’s protocol. Finally, coverslips were mounted with the Duolink *in situ* mounting media with DAPI (Sigma) on glass slides (Menzel-Gläser) and sealed. Samples were visualised on a SP5 inverted confocal microscope (Leica) using a 63X objective. Longitudinal Z-series were acquired with 0.5 µm step sizes. Image analysis was performed using the GIANI plug-in in Fiji (described in [88]). The number of foci and their distance to the edge of the DAPI staining, were quantified taking into account the 3D cell volume reflected by the longitudinal Z-series.

### Statistics

Statistical analyses were caried out using GraphPad Prism 9 software. Differences between conditions was estimated by one-way ANOVA complemented with Turkey’s *post hoc* test (*, P < 0.05; **, P < 0.01; ***, P < 0.001; ****, P < 0.0001).

## ACKNOWLEDGEMENTS

We thank David J Barry (Advanced Light Microscopy, The Francis Crick Institute) for help with microscopy data analysis. We are grateful to Joe Brock (Communications, The Francis Crick Institute) for helping design Fig 9. We acknowledge Jonathan Stoye for helpful discussions. For the purpose of Open Access, the authors have applied a CC BY public copyright licence to any Author Accepted Manuscript version arising from this submission.

---

**Supplementary table 1. Primers used to make CA mutations by site-directed mutagenesis.** Table listing forward and reverse primers used to introduce cysteine mutations on CA by site-directed mutagenesis.

**Supplementary figure 1. VLP production and Gag expression.** (A) GFP-reporter gene-expressing HIV-1 WT and mutant VLP were produced by transient transfection of 293T cells and the VLP titres in the cell supernatants were calculated by measuring RT activity using a modified RT ELISA. The bar chart shows the RT activity of the mutants relative to WT. Points indicate individual biological repeats and lines show the mean ± SEM. Colour coding is as in Fig 1. (B) Immunoblot of transfected 293T producer cell lysates probed with an anti-HIV-CA antibody showing expression of WT and mutant Gag proteins from CA mutants A14C/E45C, W184A/M185A and A14C/E45C/W184A/M185A. The blot was imaged using a LiCor Odyssey CLx imager. (C) Immunoblot of transfected 293T producer cell lysates probed with anti-HIV-CA antibody showing expression of WT and mutant Gag proteins from CA mutants A42C/T54C, Q63C/Y169C, E180C, V181C, L151C/L169C and K203C/A217C. The blot was imaged by exposure to X-ray film.

**Supplementary figure 2. Effect of CA mutations on late reverse transcription.** 293T cells were synchronously infected with equivalent RT units of WT or mutant VLP. Cells were harvested and DNA extracted and analysed for viral late cDNA products (second strand) by qPCR. (A) Bar chart shows the levels of second strand cDNA at 6 h post infection relative to WT infection. (B) Bar chart shows the levels of second strand cDNA at 24 h (left y-axis) and infectivity at 72 h (right y-axis) compared to WT VLP for each mutant. Individual points represent biological repeats and lines indicate the mean +/-SEM. (H) Bar chart shows the ratio of relative levels of second strand cDNA to infectivity, from (B). Dashed line indicates a ratio of 3. Bars are colour coded according to the lattice interface at which the cysteines have been introduced, as in fig 1. Hyper-stable mutants are indicated with black arrow heads.

**Supplementary figure 3. CPSF6 and SC35 staining during infection.** HeLa cells were synchronously infected with equal RT units of WT, A14C/E45C, E180C or M68C/E212C VLP and fixed at 16hpi. Cells were incubated with primary antibodies against CPSF6 and SC35 followed by specific secondary antibodies conjugated to Alexa Fluor fluorophores. The figure shows representative images of CPSF6 and SC35 staining at a lower magnification (63X) than in Fig 8A. The scale bar represents 20 µm.

